# Enhanced production of taxadiene in *Saccharomyces cerevisiae*

**DOI:** 10.1101/2020.06.08.139600

**Authors:** Behnaz Nowrouzi, Rachel Li, Laura E. Walls, Leopold d’Espaux, Koray Malci, Liang Lungang, Nestor Jonguitud Borrego, Albert I. Lerma Escalera, Jose R. Morones-Ramirez, Jay D. Keasling, Leonardo Rios Solis

## Abstract

Cost-effective production of the highly effective anti-cancer drug, paclitaxel (Taxol®), remains limited despite growing global demands. Low yields of the critical taxadiene precursor remains a key bottleneck in microbial production. In this study, the key challenge of poor taxadiene synthase (*TASY*) solubility in *S. cerevisiae* was revealed, and the strains were strategically engineered to relieve this bottleneck. Multi-copy chromosomal integration of *TASY* harbouring a selection of fusion solubility tags improved taxadiene titres 22-fold, up to 57 ± 3 mg/L at 30 °C at shake flask scale. The scalability of the process was highlighted through achieving similar titres during scale up to 25 mL and 250 mL in shake flask and bioreactor cultivations, respectively. Maximum taxadiene titres of 129 ± 15 mg/L and 119 mg/L were achieved through shake flask and bioreactor cultivation, respectively, of the optimal strain at a reduced temperature of 20 °C. The results highlight the positive effect of coupling molecular biology tools with bioprocess variable optimisation on synthetic pathway development.

**Highlights:**

- Maximum taxadiene titre of 129 ± 15 mg/L in *Saccharomyces cerevisiae* at 20 °C
- Integrating fusion protein tagged-taxadiene synthase improved taxadiene titre.
- Consistent taxadiene titres were achieved at the micro-and mini-bioreactor scales.

## Introduction

The highly complex diterpenoid drug Paclitaxel (Taxol^™^) first gained FDA approval in 1992 for the treatment of ovarian cancer and has since proven efficacious against a wide range of additional diseases (McElroy and Jennewein, 2018). Direct extraction from its natural source, the bark of Pacific yew (*Taxus brevifolia*), is both destructive and extremely low-yielding. As a result, paclitaxel is currently produced predominantly by semi-synthesis, involving the chemical modification of late precursors extracted from plant cell culture. However, as such methods are high in cost and have limited scalability, the development of a more sustainable source is critical to meet growing global demands (Frense, 2007). One potential solution involves the heterologous expression of the biosynthetic pathway in microbial cell factories. The first committed step in the paclitaxel biosynthetic pathway is the cyclisation of the diterpenoid intermediate, geranylgeranyl diphosphate (*GGPP*) by taxadiene synthase (*TASY*), yielding taxa-4(5),11(12)-diene (taxadiene) as shown in Figure 1.

**Figure 1:**
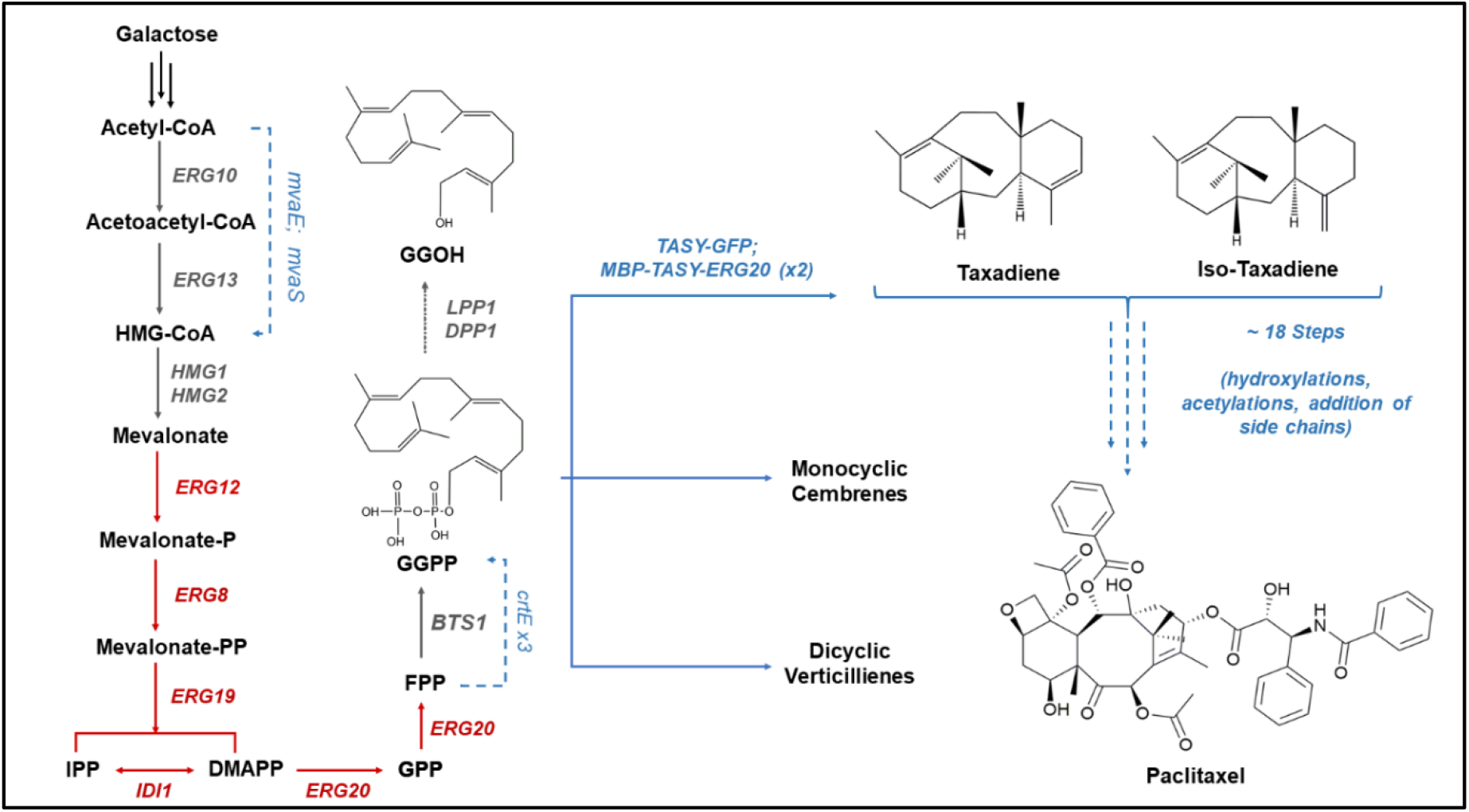
Engineered taxadiene biosynthetic pathway in *S. cerevisiae*. Genes highlighted in red represent native genes which have been overexpressed through the integration of one additional copy. Genes highlighted in blue are exogenous genes which were heterologously expressed. Hydroxymethylglutaryl-CoA synthase (*mvaS*) and Acetyl-CoA acetyltransferase (*mvaE*) from *Enterococcus faecalis.* Geranylgeranyl diphosphate synthase (*crtE*) from *Xanthophyllomyces dendrorhous* and Taxadiene synthase (*TASY*) from *Taxus cuspidata*.

TASY is comprised of three alpha-helical domains; the active site is located within the C-terminal catalytic domain, where GGPP binds and is activated via a cluster of three Mg^2+^ ions (Köksal *et al.*, 2011). TASY enzyme activity has been found to be relatively low compared to other terpene synthases, with a 70-fold lower turnover rate than that of a plant abietadiene synthase with high sequence homology (Jin *et al.*, 2005). In addition, the volume of the TASY active site is larger than that of taxadiene, contributing to previously observed enzyme promiscuity (Soliman and Tang, 2015).

The reconstitution of this enzymatic step has been successfully achieved in both *E. coli* (Huang *et al.*, 1998; Ajikumar *et al.*, 2010) and *S. cerevisiae* (Engels, Dahm and Jennewein, 2008; Reider Apel *et al.*, 2017). Through the adoption of a novel multivariate modular approach, Ajikumar *et al*. (2010) improved heterologous taxadiene titres to 300 and 1020 mg/L in *E. coli* shake-flask and fed-batch bioreactor cultivations, respectively. However, when the authors expressed the subsequent enzyme, taxadien-5α-hydroxylase, which is a membrane bound cytochrome P450, a 10-fold reduction in total taxane titre was observed. Membrane bound cytochrome P450s like taxadien-5α-hydroxylase are estimated to comprise around half of the 19 enzymatic steps in the paclitaxel biosynthetic pathway (Kaspera and Croteau, 2006). As the overexpression of such membrane bound enzymes is greatly hindered in *E. coli* (Biggs *et al*., 2016), the construction of the remainder of the pathway is likely to be very challenging in this bacterial host.

The eukaryotic host, *S. cerevisiae*, on the other hand, possesses the necessary biosynthetic machinery for the expression of such enzymes, including translocation through the endoplasmic reticulum and a native electron transfer machinery (Delic *et al.*, 2013). As in *E. coli*, early attempts to express *TASY* in *S. cerevisiae* were hindered by GGPP availability with a titre of just 122 μg/L in the wild type strain, which was insufficient for taxadiene synthesis (Engels, Dahm and Jennewein, 2008). DeJong *et al*. (2006) simultaneously incorporated GGPP synthase (*GGPPS*) and *TASY* from *Taxus* sp. into *S. cerevisiae*, leading to a taxadiene titre of 1 mg/L. Through the heterologous expression of a *Sulfolobus acidocaldarius* geranylgeranyl diphosphate synthase (*GDS*), *TASY*, and a truncated HMG-CoA, taxadiene titres were later improved to 8.7 mg/L (Engels, Dahm and Jennewein, 2008). A subsequent study focussed on the careful selection of promoters, integration locus, and solubility tags, leading to the highest reported taxadiene titre of 20 mg/L in yeast (Reider Apel *et al*., 2017).

Although taxadiene has been found to be the major product of TASY, with yields over 77%, around 5-13% of the total taxane product has been found to be the isomer taxa-4(20),11(12)-diene (iso-taxadiene) (Williams *et al.*, 2000; Huang *et al.*, 2001; Sagwan-Barkdoll and Anterola, 2018). Small quantities of a product tentatively identified as verticillene and an additional taxadiene isomer (taxa-3(4),11(12)-diene) have also been detected in *E. coli* and *Nicotiana benthamiana* (Sagwan-Barkdoll and Anterola, 2018; Li *et al.*, 2019).

The metabolic pathway of paclitaxel is highly complex and development of an alternative recombinant production route remains in the preliminary stages. Despite this, substantial advancements in synthetic biology have been achieved recently (Hsu, Lander and Zhang, 2014).

Through the application of such tools, there is great potential to accelerate the development of a microbial paclitaxel biosynthetic pathway. This study focussed on the optimisation of TASY enzyme performance in *S. cerevisiae* to alleviate a key early pathway bottleneck. The effect of a number of factors including TASY truncation length, selected promoter and chromosomal gene copy number on pathway expression were evaluated. Cultivation conditions such as culture temperature and exogenous cofactor availability were also considered.

## 2. Materials and methods

### 2.1. Yeast strains and media

The parent *S. cerevisiae* strain used for episomal expression and integration studies was GTy116 (MATa, *leu2-3, 112::HIS3MX6-GAL1p-ERG19/GAL10p-ERG8; ura3-52::URA3-GAL 1p-MvaSA110G/GAL10p-MvaE* (codon-optimised); *his3Δ1::hphMX4-GAL1p-ERG12/GAL10p-IDI1;trp1-289::TRP1_pGAL1-CrtE(X.den)/GAL10p-ERG20; YPRCdelta15::NatMX-GAL1p-CrtE(opt)/GAL10p-CrtE*) described previously by Reider Apel *et al*. (2017), originating from CEN.PK2-1C (EUROSCARF collection). The *URA3* marker of this strain was further restored to give mGty116. All chemicals and reagents were sourced from Sigma-Aldrich at the highest available purity unless otherwise stated. Episomal expression systems made use of synthetic defined medium minus uracil (CSM-Ura and CSM-Leu, Sunrise Science Products) depending on the selection marker used. These were supplemented with 2% *(w/v)* glucose (SDD-Leu), 2% *(w/v)* galactose (SDG-Leu) or a 1.8% *(w/v)* galactose, 0.2% *(w/v)* glucose mixture (SDGD-Leu). For cultivation of strains harbouring chromosomally integrated genes, a medium containing yeast extract (*1%(w/v)*) and peptone (2 %*(w/v)*), supplemented with 2 % *(w/v)* glucose (YPD), galactose (YPG) or 1.8% *(w/v)* galactose and 0.2% *(w/v)* glucose mixture (YPGD) were used.

### 2.2. Yeast transformation and strain construction

Episomal expression was achieved through transforming *S. cerevisiae* with high copy 2-micron plasmids harbouring *TASY* and a *LEU2* selection marker (Supplementary Table 1). The standard LiAc/SS carrier DNA/PEG method was used for all transformations (Gietz and Schiestl, 2007). Chromosomal integration was performed using a cloning free, Cas9-mediated homologous recombination method (Reider Apel *et al.*, 2017). High copy number, Cas9-sgRNA 2-micron plasmids harbouring a *URA3* selection marker derived from pRS426 were used. The gene cassettes were designed using an online tool, CASdesigner. Chromosomal integration at the target site was confirmed by colony PCR and Sanger sequencing (Genewiz Inc., USA; Edinburgh Genomics, UK). Plasmid curing was subsequently performed on successful colonies through sequential culture on YPD agar medium until no growth was observed on concurrent SDD-Leu agar plates.

The strains used in this study are summarised in Table 1 and plasmid, primer and tag sequences are tabulated in Supplementary Tables 1-3. All DNA sequences were synthesised by IDT (Integrated DNA Technologies, Inc.). A *TASY* sequence from *Taxus cuspidata* was codon optimised for expression in *S. cerevisiae*. DNA amplification was performed using Phusion Flash High-Fidelity PCR Master Mix (Thermo Fisher Scientific). Promoter and terminator sequences native to the yeast genome were selected (Lee *et al.*, 2015).

**Table 1:**
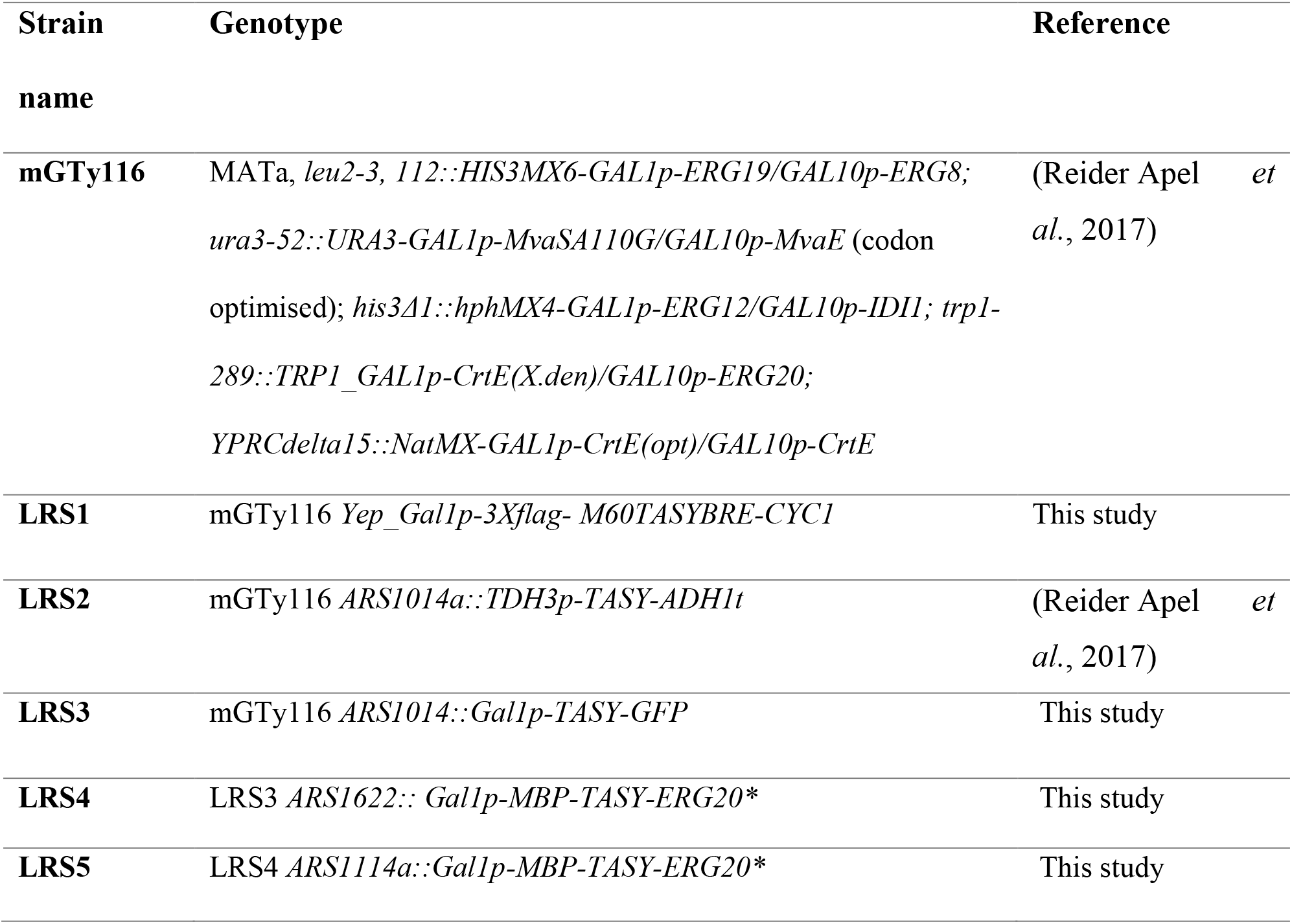
Yeast strains used in this study.

### 2.3. Optimisation of taxadiene production at microscale

Preliminary microscale episomal expression studies involved 5 ml cultures of the mGTy116 strain harbouring selected high-copy 2-micron plasmids as shown in Supplementary Table 1. Inocula were prepared through transferring a single yeast colony to 5 ml of SDD-Leu medium, followed by incubation overnight at 30 °C. Aliquots of the inoculum were diluted with SDGD-Leu medium to give an initial OD_600_ = 0.5. The yeast cultures were incubated at 20 or 30 °C and 200 rpm for 96 hours. To minimise air-stripping of the volatile terpene products, a biphasic liquid-liquid system was used through the addition of a 10 % *(v/v)* dodecane overlay after 24 hours of cultivation. At the end of the cultivation, the resulting cultures were centrifuged and the organic dodecane layer was extracted for GC-MS analysis. Microscale culture of the strains harbouring chromosomally integrated genes was done in a similar way as for episomal expression, except that YPD and YPGD media were used for the inoculum and production cultures, respectively.

### 2.4. Shake flask cultivations

Single LRS5 colonies were transferred from YPD agar to 5 mL YPD medium and incubated at 30 °C and 250 rpm overnight. Aliquots of these cultures were then used to inoculate 20 mL YPG in 250 mL shake flasks to an OD_600_ = 1. A 20% *(v/v)* dodecane overlay was used giving a total working volume of 25 mL. The resulting cultures were incubated at 20 or 30 °C and 250 rpm for 72 hours. At the end of the cultivation, the dodecane overlay was extracted for GC-MS analysis. The final biomass was measured at an optical density of 600 nm.

### 2.5. Batch culture in the bioreactor

Cultivations were conducted in MiniBio 500 mL bioreactors (Applikon Biotechnology, The Netherlands) with a working volume of 250 mL. Pre-inocula were prepared by incubating cells in 5 mL of YPD for eight hours. The resulting cultures were used to inoculate secondary 10-mL inocula to an OD_600_ = 1, which were subsequently incubated at 30 °C and 200 rpm overnight. An aliquot of an inoculum culture was then diluted with YPG to give a 200 mL culture with an initial OD_600_ = 1. To prevent excess foam production, polypropylene glycol P2000 (Fisher Scientific, UK) was added to a concentration of 0.01 % *(v/v)* and a Rushton turbine was placed at the medium-air interface. A 20 % *(v/v)* dodecane (Fisher Scientific, UK) overlay was also added to minimise product loss due to air stripping. During cultivation, the temperature, dissolved oxygen and pH were monitored online. Biomass was measured through manual sampling twice daily. The adaptive my-Control system (Applikon Biotechnology, The Netherlands) was used to control process parameters. Setpoints of 30 % of the saturation and 30 °C were applied for dissolved oxygen and temperature, respectively. The culture pH was controlled to a setpoint of six through automatic addition of 1M NaOH. Samples were taken twice daily for analysis of taxane and biomass concentration.

### 2.6. Diterpene analysis and quantification

The dodecane overlay was analyzed by GC-MS using Trace 1300 GC (ThermoFisher Scientific), equipped with TG-SQC column (15 m × 0.25 mm × 0.25 μm). The mass spectra in the range of 50-650 m/z was recorded on a Thermo Scientific ISQ Series single quadrupole mass spectrometer using EI ionization mode and scan time of 0.204 s. The GC temperature programme began at 120 °C (3 min) and was then raised to 250 °C at a rate of 20 °C/min with 3 min hold time. Xcalibur^™^ software (ThermoFisher Scientific, USA) was employed for data processing. Pure taxadiene (kindly supplied by the Baran Lab, The Scripps Research Institute) and geranylgeraniol (Sigma Aldrich, UK) were used as standards to identify and quantify taxadiene and GGOH, respectively. Additional taxane products were quantified relative to standard taxadiene concentrations.

### 2.7. Intracellular taxadiene quantification

Intracellular taxadiene accumulation was quantified in the 20 °C bioreactor samples. Cell lysis was achieved using a previously described protocol (Westfall *et al.*, 2012) with some modifications. In summary, 1 mL of the aqueous culture phase was centrifuged and the cell pellet was resuspended in a mixture containing 0.4 mL of yeast lysis reagent (2 % *(v/v)* Triton X100,1 % *(w/v)* SDS, 100 mM NaCl, 10 mM Tris-HCl (pH 8.0) and 1 mM EDTA (pH 8.0)) and 0.4 mL of 1M HCl for cell lysis. The resulting suspension was mixed thoroughly prior to the addition of a 1-mL dodecane overlay. The samples were subsequently incubated at 30 °C, 200 rpm for 16 hours before separation of the organic phase for GC-MS analysis.

## 3. Results and discussion

### 3.1. Optimisation of taxadiene titre using episomal *TASY* expression

Four key parameters with the potential to affect final taxadiene titre were examined using a high-copy (2-micron) plasmid in the mGty116 *S. cerevisiae* strain. Such parameters included the selected promoter, cultivation temperature, cofactor (Mg2^+^) concentration, and *TASY* sequence truncation. The results are summarised in Figure 2.

**Figure 2:**
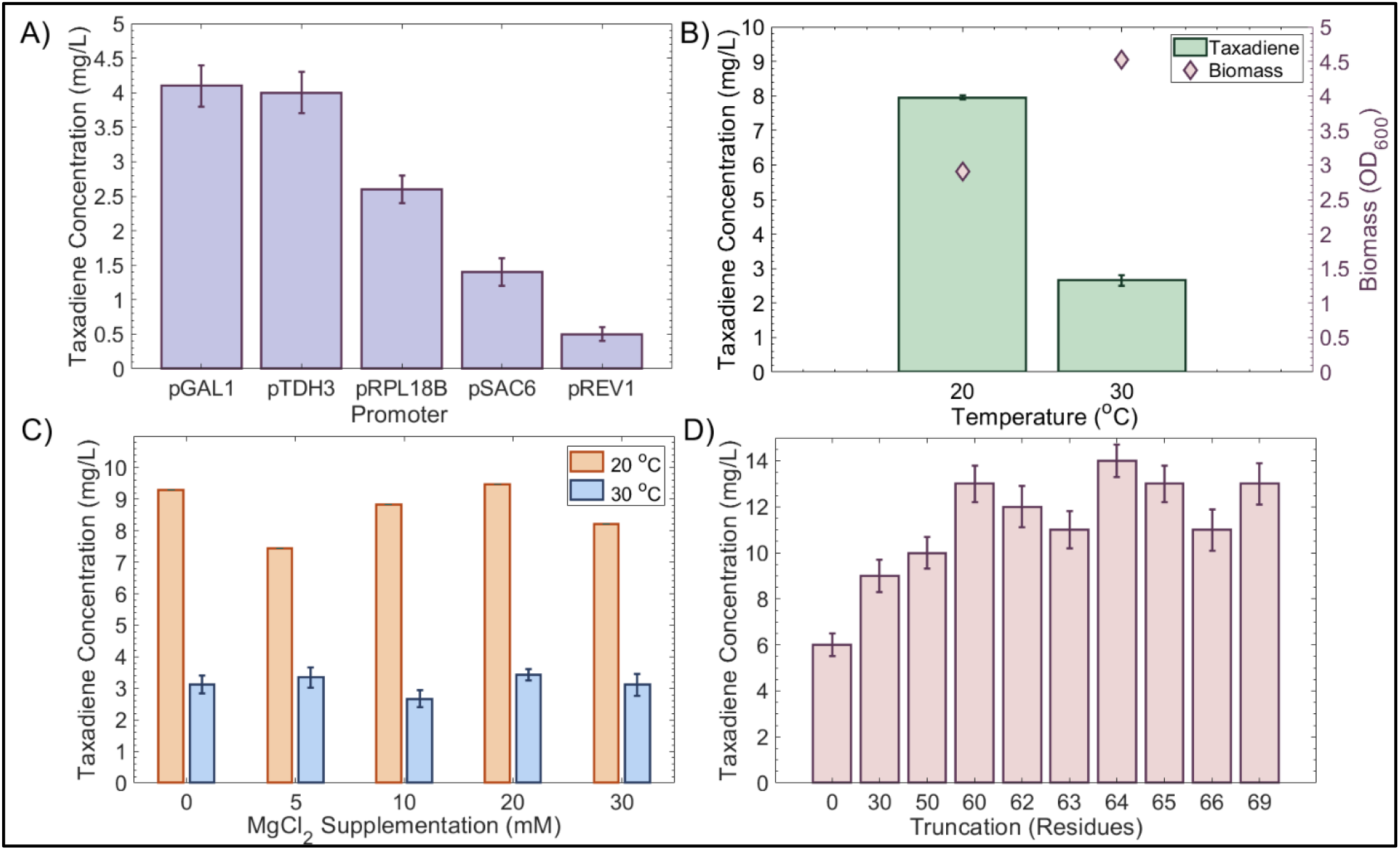
Optimisation of taxadiene titres using episomal *TASY* expression. A) Effect of promoter on taxadiene concentration produced by cells incubated at 30 °C. B) Effect of cultivation temperature on taxadiene titre. C) Effect of cofactor (Mg^2+^) availability at 20 and 30 °C. D) Effect of gene truncation length for taxadiene production at 20 °C. Values shown are mean ± standard deviation for 96-hour triplicate cultivations.

In this study, the performance of four constitutive promoters of increasing strength, pREV1, pSAC6, pRP118BP and pTDH3 (Lee *et al.*, 2015) were compared to that of the strong inducible GAL1 promoter in *S. cerevisiae*, grown in SDGD-Leu media (Figure 2A). Performance of the highest strength constitutive TDH3 and inducible GAL1 promoters was highly similar with titres of 4.1 ± 0.3 and 4.0 ± 0.3 mg/L, respectively. Decreasing promoter strength did not improve taxadiene titres, which were 1.6, 2.9 and 8.2-fold lower for the weaker, RPL118, SAC6 and REV1 promoters, respectively. As changing the promoter from the inducible GAL1 promoter resulted in no further improvement in taxadiene titre (Figure 2A), pGAL1 was selected for subsequent studies.

The growth temperature was found to have an important effect on taxadiene titre as shown in Figure 2B. Reducing the cultivation temperature from 30 °C to 20 °C resulted in a 3.0-fold increase in final taxadiene titre to 8 ± 0.l mg/L. However, at the lower temperature biomass accumulation was also reduced, with a final OD_600nm_ of 2.90, compared to 4.5 at 30 °C. This was in agreement with previous results where heterologous expression of *TASY* in *E. coli* (Boghigian *et al.*, 2012) and *Bacillus subtilis* (Abdallah *et al.*, 2019) has also been found to be sensitive to culture temperature. Productivity analysis over a broad temperature range (12-37 °C) revealed a temperature optimum of 22 °C for both species (Boghigian *et al.*, 2012). In addition, the highest reported titre of 1020 mg/L was achieved in *E. coli* cultures at a temperature of 22 ^°^C (Ajikumar *et al.*, 2010). Similarly, a recent study by Abdallah *et al*. (2019) showed that *TASY* expression in *B. subtilis* was best at 20 °C, as opposed to 30 °C or 37 °C.

TASY relies on the metal ion co-factor, Mg^2+^, for activation and substrate orientation within the active site (Schrepfer *et al.*, 2016). In a recent study by Tashiro *et al*. (2018), MgCl_2_ supplementation was found to improve activity of an alternative diterpene synthase, pinene synthase, up to 20 mM in *E. coli*. As the concentration of Mg^2+^ in the SDGD-Leu medium was around 0.8 mM, it was hypothesised to be rate limiting. However, supplementation of the cultivation medium with additional MgCl_2_ did not significantly improve taxadiene titres at 20 or 30 °C in this study (Figure 2C). Consequently, additional MgCl_2_ supplementation was not deemed necessary for subsequent experiments.

The native *TASY* gene encodes 862 amino acids, including a putative N-terminal sequence of ~137 residues which is cleaved upon maturation in plastids (Köksal *et al.*, 2011). Removal of this sequence was found to reduce inclusion body formation and increase active protein production in *E. coli* (Williams *et al.*, 2000; Köksal *et al.*, 2011). In order to determine the optimal protein length for TASY expression in *S. cerevisiae*, a range of truncation lengths were tested (Figure 2D). Increasing truncation length up to 60 residues improved taxadiene production from 6.0 ± 0.5 to 13.0 ± 0.8 mg/L as shown in Figure 2D. In variants harbouring longer truncations, no further improvement in taxadiene titre was observed. The *TASY* variant with a 60-residue truncation was therefore selected for subsequent experiments. These results were in agreement with previous works where truncations of 60 or 79 residues yielded active protein and reduced inclusion body formation, whilst truncations of 93 or more residues produced inactive protein in *E. coli* (Williams *et al.*, 2000).

### 3.2. Chromosomal integration of *TASY*

#### 3.2.1. Comparing episomal and chromosomal expression

Plasmid-based systems rely on the use of expensive selective media, limiting their industrial relevance. In a previous study, a single copy of the *TASY* gene was chromosomally integrated into the mGty116 strain. Cultivation of the resulting LRS2 (pTDH3-*TASY*-tADH1) strain at 30 °C resulted in a taxadiene titre of 2.6 ± 0.4 mg/L (Reider Apel *et al.*, 2017). Interestingly, this titre was comparable to the 2.7 ± 0.2 mg/L achieved using the high-copy 2-micron plasmid system in this study (Figure 2B). The effect of stable chromosomal *TASY* integration in yeast was therefore investigated in the mGty116 strain of this study. In order to visualise functional TASY expression, *TASY* was tagged with a C-terminus GFP reporter gene (strain LRS3). Cultivation of this strain yielded 12 ± 1 mg/L of taxadiene at 30 °C, 4.7-fold higher than that of the LRS2 strain. This was also comparable to the optimal plasmid-based titre of 13 ± 1 mg/L achieved at the lower temperature of 20 °C (Figure 2D). This suggested that although chromosomal integration improved taxadiene titre in comparison to episomal expression, solubility remained a significant bottleneck.

#### 3.2.2. Effect of gene copy number and solubility tags

Fusion tags are often used to improve protein expression and solubility (Kosobokova, Skrypnik and Kosorukov, 2016). A recent study using a range of tags, revealed that maltose binding protein (MBP), had the greatest positive impact on TASY solubility and activity in *S. cerevisiae* (Reider Apel *et al*., 2017). Prenyltransferase-terpene synthase fusion proteins have also been found to greatly improve terpene production (Ignea *et al.*, 2015; Deng *et al.*, 2016). The prenyltransferase, farnesyl diphosphate synthase, encoded by *ERG20* catalyses the formation of the geranyl diphosphate (GPP) and farnesyl diphosphate (FPP) metabolites, two key enzymatic steps in the mevalonate pathway (Figure 1). ERG20 has been reported to significantly improve isoprenoid concentration when used as a fusion protein with terpene synthases, potentially promoting enzyme solubility as well as increasing the pool of the isoprenoid precursor farnesyl diphosphate (Ignea *et al.*, 2015; Wong *et al.*, 2018). Deng *et al*. (2016) showed that the fusion of a mutant variant of *ERG20* to (*S*)-linalool synthase improved the productivity by around 70% compared to independent integration of the two genes in *S. cerevisiae*. Hence, a cassette containing an N-terminal yeast codon-optimised *MBP* tagged *TASY-ERG20*\* (F96C; Ignea *et al.*, 2015) fusion gene was developed with the dual aim of increasing precursor availability and improving TASY solubility. This cassette was chromosomally integrated into locus ARS1014a of LRS3 to yield strain LRS4. The integration of a second copy of this dual-tagged *TASY* in locus ARS1622b resulted in strain LRS5 (1x *TASY-GFP*, 2x *MBP-TASY-ERG20*\*, Figure 3).

**Figure 3:**
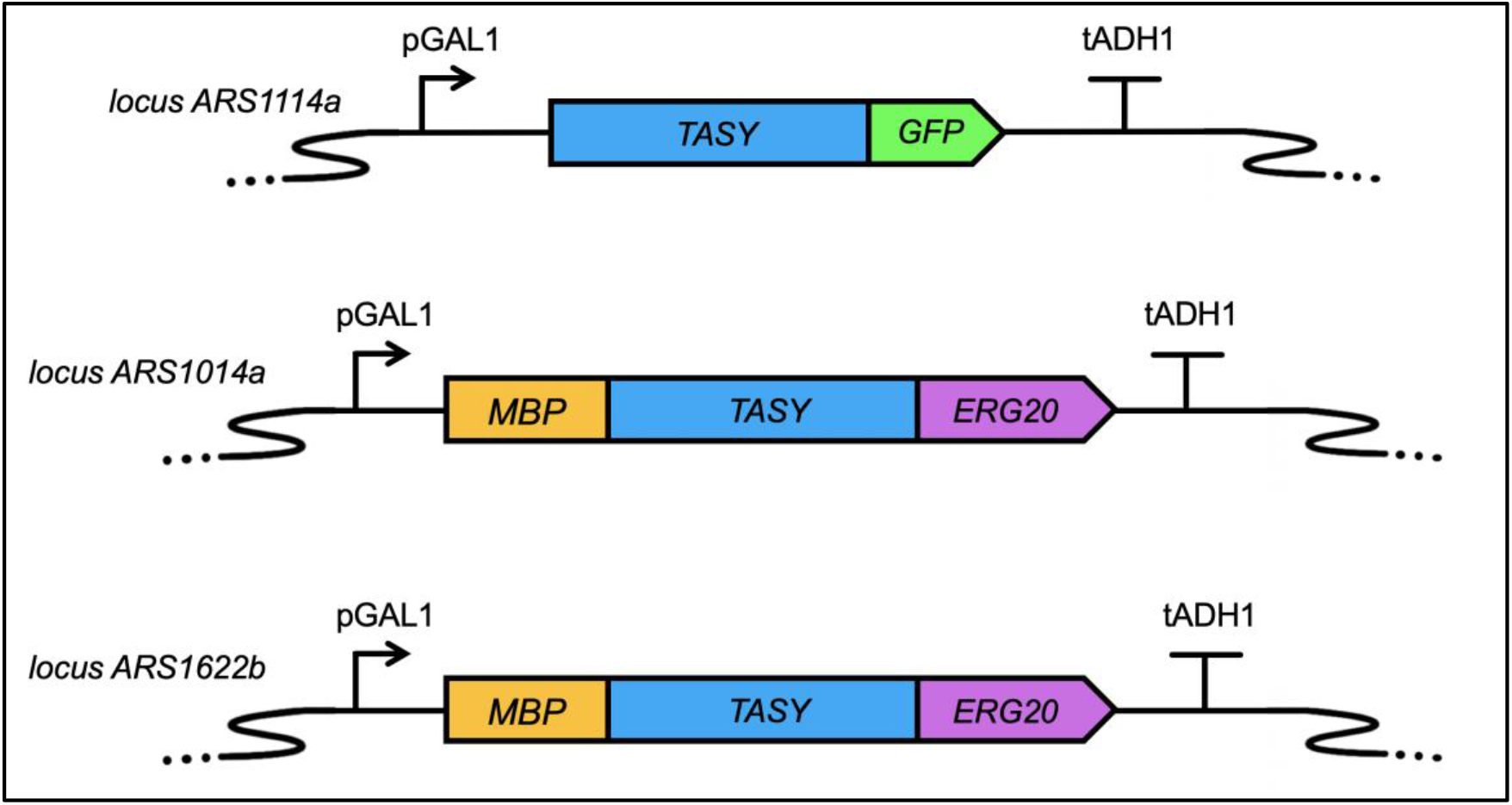
Chromosomally integrated *TASY* genetic constructs. From top to bottom are *TASY* cassettes integrated at loci: ARS1114a, ARS1014a, and ARS1622b, respectively. The genetic designs were visualized by SBOLDesigner 2 (Zhang *et al.*, 2017).

Cultivation of the newly constructed LRS4 strain expressing two copies of *TASY* resulted in an over three-fold improvement in taxadiene titre to 40 ± 3 mg/L at microscale. Through cultivation of the subsequent LRS5 strain, taxadiene titres were further enhanced to 57 ± 3 mg/L at 30 °C, the highest titre in yeast reported to date at microscale level (Figure 4).

**Figure 4:**
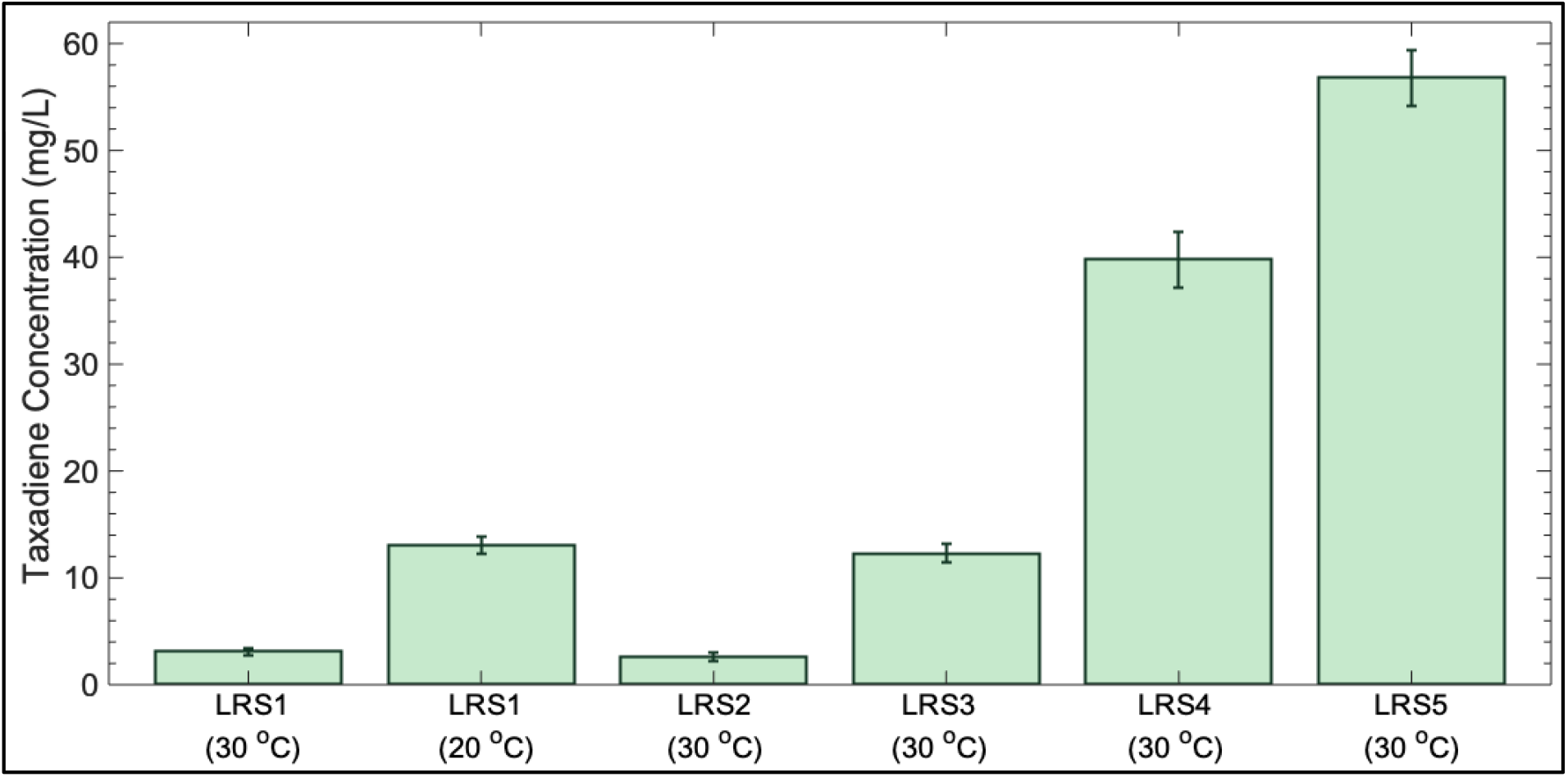
Summary of taxadiene titers microscale optimisation study. LRS1 indicates titers resulting from episomal *TASY* expression using the high copy 2-micron plasmid (LRS1) at 30 and 20 ^°^C, respectively. The other titers are from strains with chromosomally integrated *TASY* variants: LRS2 (TDH3p-*TASY*-ADH1t), LRS3 (*TASY*-*GFP*), LRS4 (*TASY*-*GFP*; *MBP*-*TASY*-*ERG20*\*) and LRS5 (*TASY-GFP; MBP-TASY-ERG20*; MBP-TASY-ERG20*\*), respectively, grown at 30 °C.

Through effective copy number optimisation and the use of MBP and GFP tags along with an ERG20 fusion, taxadiene production was enhanced up to 22-fold in *S. cerevisiae* (Figure 4). This was likely the result of improved protein expression and solubility, as well as precursor availability. Nevertheless, fluorescent visualisation still showed substantial spotted subcellular localization (Supplementary Figure 1), which was consistent with a previous study highlighting the poor solubility of TASY (Reider Apel *et al.*, 2017).

### 3.3. Taxadiene production scale-up

#### 3.3.1. Minor product characterisation at increased scale under different temperatures

In order to investigate process scalability, the optimised LRS5 strain was subsequently cultivated in 250-mL shake flasks (25 ml working volume). The effect of temperature on chromosomal *TASY* expression by the optimised strain was also assessed through cultivation at both 20 °C and 30 °C. The results of this experiment are summarised in Figure 5.

**Figure 5:**
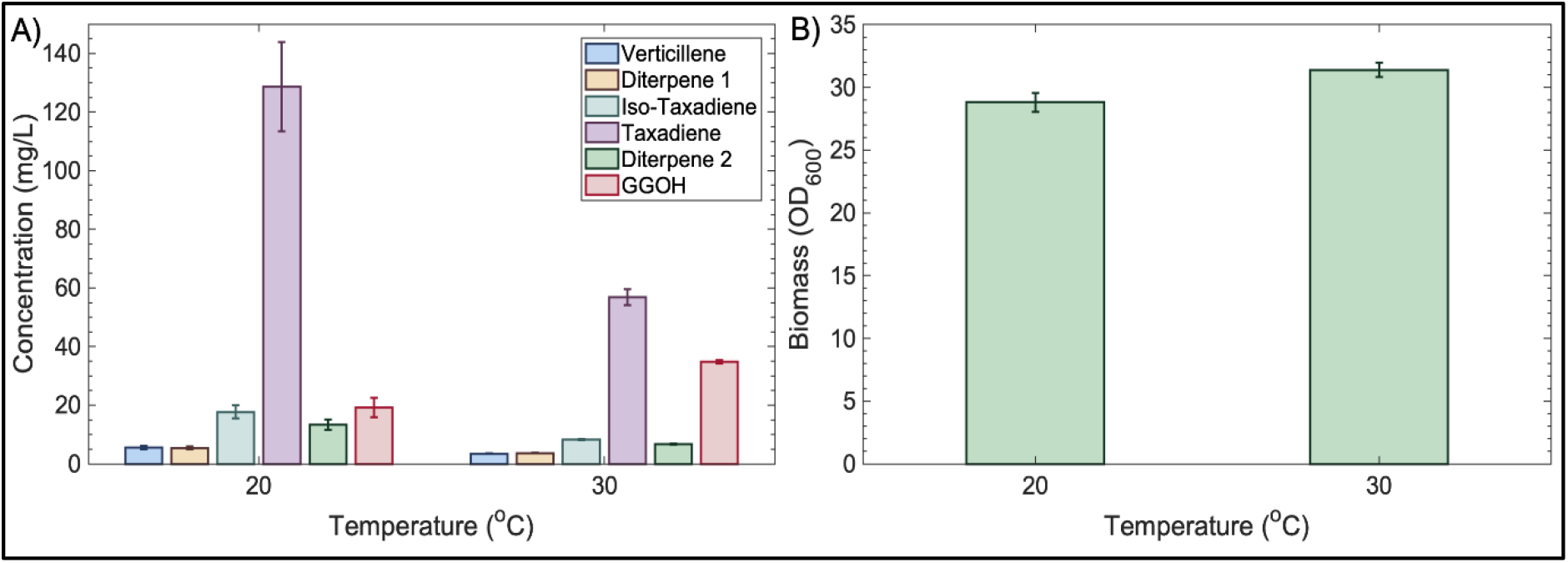
Effect of temperature on LRS5 performance in shake flasks. LRS5 was cultivated in 250 mL shake flasks in YPG media at 20 or 30 °C. Taxane accumulation (A) and yeast growth (B) were evaluated after 72 hours of cultivation. Error bars represent ± standard deviation for triplicate cultivations.

The final taxadiene titre at 20 °C was 129 ±15 mg/L, representing an almost 6.5-fold improvement over the highest reported titre (Reider Apel *et al.*, 2017). The final titre at 30 °C was 56.9 ± 2.7 mg/L, which was comparable to the 57 ± 3 mg/L obtained in the smaller scale cultivations (Figure 6). This represented a 2.3-fold lower taxadiene titre at 30 °C compared to 20 °C, indicating that despite the addition of solubility tags, further optimisation is likely to be necessary at higher growth temperatures. Interestingly, biomass accumulation was very similar for the strain at the two temperatures, with final OD_600_ values of 29 ± 1 and 31 ± 1 at 20 and 30 °C, respectively (Figure 5B).

**Figure 6:**
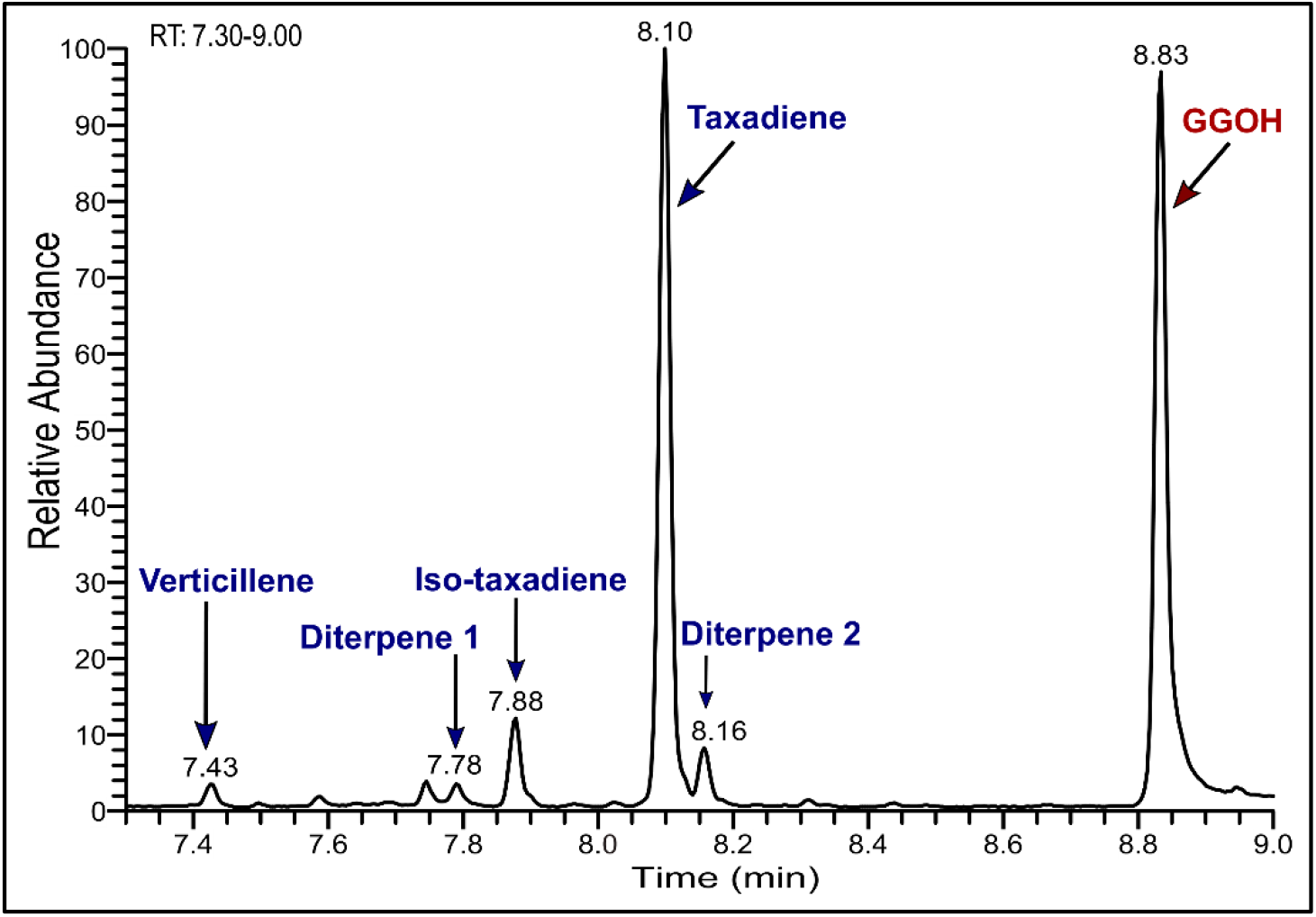
LRS5 gas chromatogram. Results show products produced by LRS5 during the 30 °C shake flask cultivation.

Although taxadiene was the major product under all conditions studied, additional side products were also generated. At this scale, further characterisation of such side products was performed through a detailed analysis of gas chromatography (Figure 6) and mass spectrometry data (Supplementary Figure 2).

In addition to taxadiene, which was eluted at 8.10 minutes (Supplementary Figure 2D), a further five diterpene products were detected as shown in Figure 6. The mass spectra of the peaks at 7.43 and 7.88 minutes (Supplementary Figures 2A and C) showed a high degree of similarity to compounds previously identified as verticillene and the structural taxadiene isomer, taxa-4(20),11(12)-diene (iso-taxadiene), respectively (Williams *et al.*, 2000; Sagwan-Barkdoll and Anterola, 2018). An additional two diterpene compounds, Diterpene 1 and Diterpene 2 (Supplementary Figures 2B and E) were observed at 7.78 and 8.16 minutes, respectively. The mass spectrum of diterpene 1 closely resembled cembrene-type diterpene (cembrene A) produced by neocembrene synthase in *S. cerevisiae* (Kirby *et al.*, 2010). The cyclisation of GGPP to taxadiene involves production of a cembrenyl intermediate (van Rijn, Escorcia and Thiel, 2019). The relatively high degree of similarity suggests the compound observed in this study could potentially be a side product of this cyclisation step. However, further characterisation is needed to confirm this. The mass spectrum and elution order of diterpene 2 was very similar to that of a novel diterpene product of *TASY* expression in *Nicotiana benthamiana* (Li *et al.*, 2019). Production of geranylgeraniol (GGOH) (8.83 minute; Supplementary Figure 2F) was also confirmed using a pure analytical standard. Although GGOH is formed naturally by *S. cerevisiae* through the degradation of GGPP by endogenous phosphatases, titres are typically too low to detect (Song *et al.*, 2017). Its overproduction here indicates that early work to overexpress the mevalonate pathway was successful, however, taxadiene titres could be improved further through pathway optimisation. In addition to the diterpene products, a number of additional potential terpenoid side-products were detected between 4.36 and 6.38 minutes (Supplementary Figures 3 and 4).

#### 3.3.2. Scale-up using a mini-bioreactor system

A 10-fold scale-up of the LRS5 cultivation was used to assess performance under industrial conditions for taxadiene production. A 500-mL MiniBio bioreactor (Applikon, UK) was employed for this study. This system possesses the online monitoring and control capabilities of larger-scale bioreactors, allowing industrial scale cultivation conditions to be effectively mimicked whilst conserving valuable resources. The results of the bioreactor-scale runs are summarised in Figure 7.

**Figure 7:**
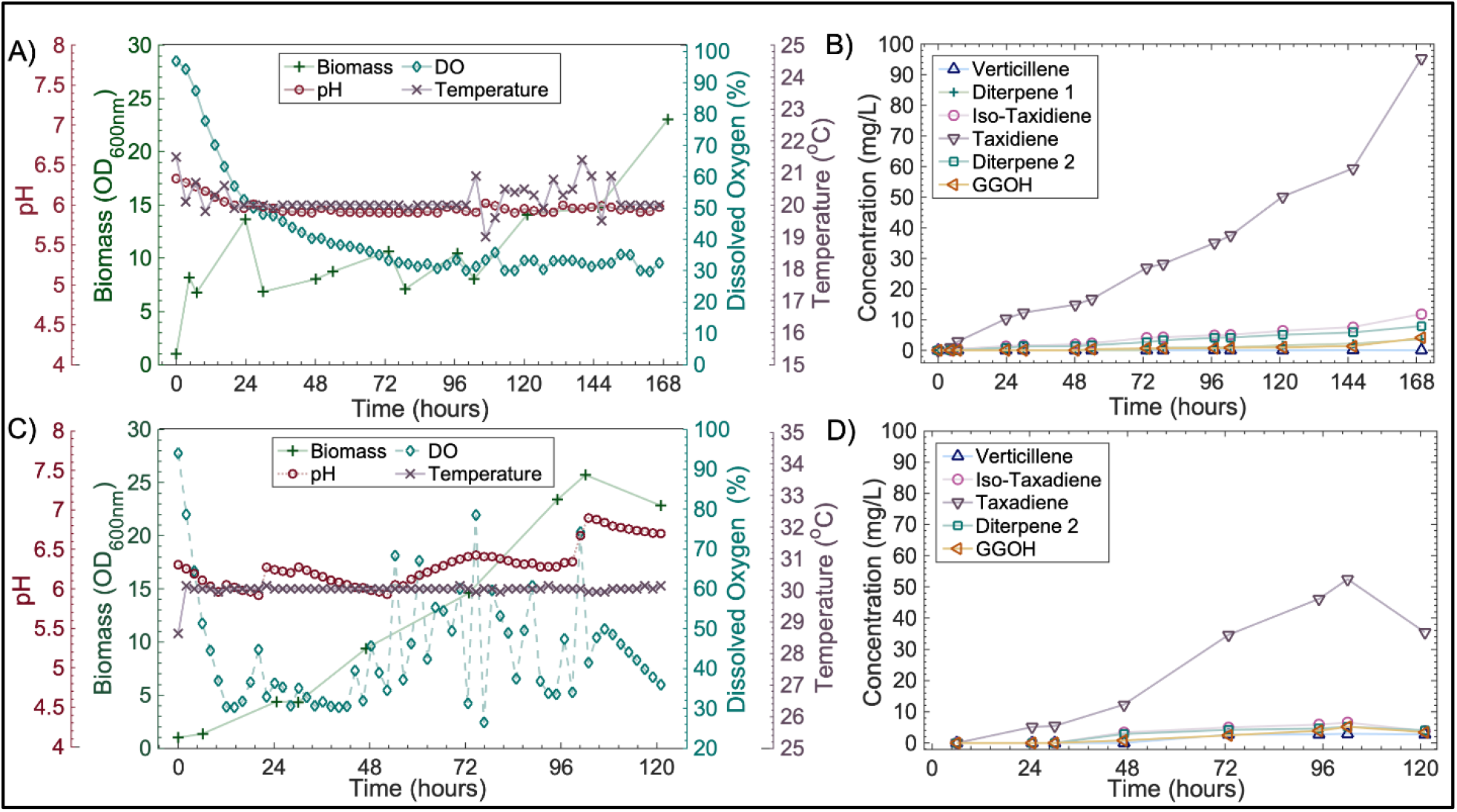
Bioreactor studies results and respective taxane concentration kinetics at 20 °C (A and B) and at 30 °C (C and D). LRS5 was cultivated in an Applikon MiniBio 500 bioreactor in yeast extract (1%), peptone (2%), galactose (2%). The pH and dissolved oxygen were monitored online and controlled to set points of 6 and 30%, respectively online.

Following 72 hours of cultivation, the OD_600_ values of the 20 and 30 °C cultivations were 10.6 and 14.6 respectively as shown in Figures 7A and C. This is significantly lower than the 29 ± 1 and 31 ± 1 obtained at shake flask scale (Figure 5B). Taxadiene titres were also lower in the bioreactor studies at 27 and 35 mg/L compared to 129 and 57 mg/L for the equivalent 20 and 30 °C shake flask cultures. Despite this, following a further 30 hours of cultivation, the stationary phase of growth was reached and a maximum taxadiene titre of 53 mg/L was attained in the 30 °C reactor. This is highly comparable to those obtained in the 72-hour microscale (Figure 6) and shake flask (Figure 5A) cultivations. A higher taxadiene titre of 109 mg/L was achieved in the 20 °C bioreactor, however, a bioreactor run time of 195 hours was required. Interestingly, production of the endogenous GGOH side product was dramatically reduced at the increased scale with maximum titres of 4 and 5 mg/L (Figures 7B and D) compared to 19 ± 3 and 35 ± 1 mg/L for the equivalent 20 and 30 °C shake flask studies (Figure 5A).

Unlike prokaryotes, eukaryotic yeast cells have evolved to store hydrophobic compounds within specialised organelles (liquid droplets) (Arhar and Natter, 2019). As a result, it was hypothesised that additional taxadiene may be present within the yeast cells. An investigation into intracellular taxane production was therefore performed on samples taken from the higher-yielding 20 °C bioreactor cultivation as summarised in Figure 8.

**Figure 8:**
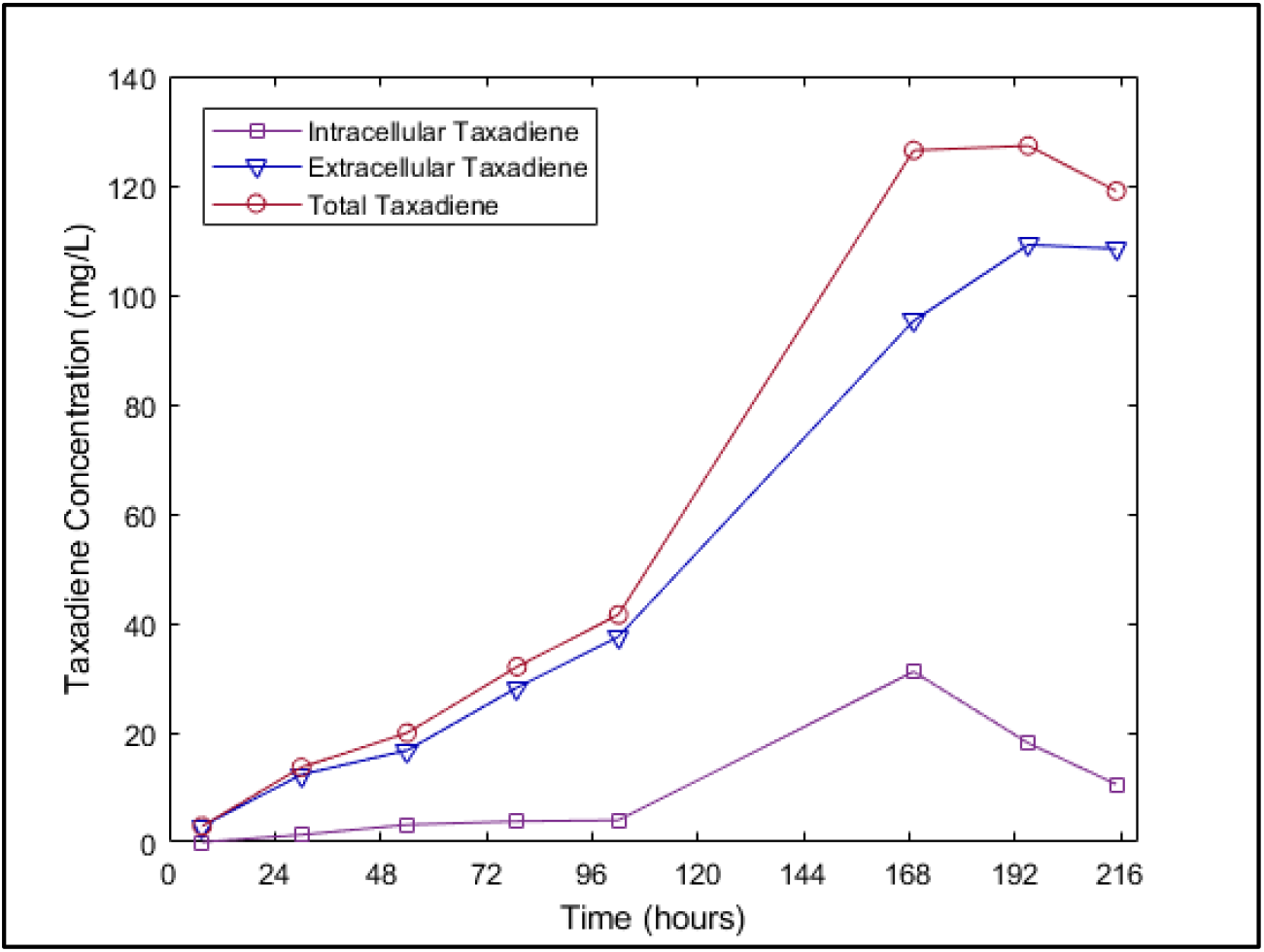
Kinetics of intracellular and total taxadiene titres in high-yielding 20 °C bioreactor cultivation.

In the first 168 hours of the cultivation, both extracellular and intracellular taxadiene accumulation increased (Figure 8). In the subsequent 48 hours a sharp decline in intracellular taxadiene from 31 to 11 mg/L was observed. This was coupled with an increase in extracellular taxadiene from 95 to 109 mg/L while the total taxadiene concentration remained similar between 119 and 127 mg/L (Figure 8). This suggested that the increase in extracellular taxadiene concentration observed in the last 48 hr was likely largely due to taxadiene secretion rather than additional taxadiene biosynthesis.

A lower cultivation temperature of 20 °C was found to improve TASY performance with higher taxadiene titres being obtained under all the conditions investigated in this study. However, the kinetic analysis revealed a strong positive correlation between biomass and product accumulation at both 20 and 30 °C (Pearson’s R = 0.916 and 0.974, respectively). As the growth rate of LRS5 decreased with temperature, longer cultivation time was required at 20 °C. This was exacerbated at larger scale, where cultivation time was tripled (Figure 7A). Although careful optimisation yielded substantial improvements in TASY productivity in this study, the discrepancies between the optimal temperature for the enzyme and host growth remain a challenge. The optimisation of copy number and protein tag combination improved taxadiene titres dramatically in this study. However, the higher titres obtained at reduced temperature indicate further optimisation is needed at the optimal growth temperature of 30 °C.

## 4. Conclusions

This study reports the successful optimisation of taxadiene biosynthesis in a *Saccharomyces cerevisiae* microbial chassis. Low expression and poor solubility of taxadiene synthase (TASY) were identified as a critical bottlenecks. This was alleviated through multi-copy chromosomal integration of *TASY* with a combination of fusion protein tags, improving taxadiene titres 22-fold to 57 ± 3 mg/L at 30 °C. TASY performance was found to be temperature-dependant, with a maximum taxadiene titre of 129 ± 15 mg/L at 20 °C. Similar titres were achieved at larger scale as well, highlighting the scalability of the bioprocess, and representing a 6.5-fold improvement on the highest literature titre.

This work highlights the benefit of multifactorial approach to biosynthetic pathway optimisation, where a combination of synthetic biology tools and bioprocessing approaches were employed. Although the current study presents significant progress in taxadiene biosynthesis, several tasks remain to improve the expression of taxadiene synthase at 30 °C to allow for further Paclitaxel biosynthetic pathway development in *S. cerevisiae*.

## 5. Acknowledgements

The authors would like to thank Professor Phil Baran’s Lab at The Scripps Research Institute, San Diego, California for providing the taxadiene standard. Thanks to Ms. Caroline Delahoyde and Mr. Martin Corcoran for their kind assistance and technical support with GC-MS analysis. This work was supported by The University of Edinburgh’s Principal Career development Scholarship, The Engineering and Physical Sciences Research Council [grant number: EP/R513209/1], YLSY Program of the Ministry of National Education of the Republic of Turkey, the Royal Society [grant number: RSG\R1\180345], University of Edinburgh Global Challenges Theme Development Fund [grant number: TDF_03] and the US National Science Foundation (Award Number 1330914).

## Abbreviations

BTS1: Geranylgeranyl diphosphate synthase
*crtE*: Geranylgeranyl diphosphate synthase
DMAPP: Dimethylallyl pyrophosphate
*ERG8*: Phosphomevalonate kinase
*ERG9*: Farnesyl-diphosphate farnesyl transferase (squalene synthase)
*ERG10*: 3-hydroxy-3-methylglutaryl-CoA (HMG-CoA) synthase
*ERG12*: Mevalonate kinase
*ERG13*: 3-hydroxy-3-methylglutaryl-CoA (HMG-CoA) synthase
*ERG19*: Mevalonate pyrophosphate decarboxylase
*ERG20*: Farnesyl pyrophosphate synthetase
FPP: Farnesyl diphosphate
GGOH: (E,E,E)-geranylgeraniol
GGPP: (E,E,E)-Geranylgeranyl diphosphate
GPP: Geranyl diphosphate
*HMG1*: 3-hydroxy-3-methylglutaryl-coenzyme A reductase 1
*HMG2*: 3-hydroxy-3-methylglutaryl-coenzyme A reductase 2
*IDI*: Isopentenyl-diphosphate delta-isomerase
*IPP*: Isopentenyl pyrophosphate
MBP: Maltose binding protein
*mvaE*: Acetyl-CoA acetyltransferase
*mvaS*: Hydroxymethylglutaryl-CoA synthase
MVA pathway: Mevalonate pathway
*TASY*: Taxadiene synthase.

