## Supplementary Information for "Enhanced production of taxadiene in *Saccharomyces cerevisiae*"

### Appendix A. Supplementary material

Supplementary Table 1: **Plasmids used in this study**

| **Plasmid name** | **Description** |
|---|---|
| pLR1 | Yep_pGAL1-3Xflag-M60TASYBRE-tCYC1 |
| pLR2 | Yep_pSV1-3Xflag-M60TASY-tCYC1 |
| pLR3 | Yep_pVAN1-3Xflag-M60TASYBRE-tCYC1 |
| pLR4 | Yep_pADH1-3Xflag-M60TASYBRE-tCYC1 |
| pLR5 | Yep_pTDH3-3Xflag-M60TASYBRE-tCYC1 |
| pL10 | Yep_pGAL1-M60TASYBRE-tCYC1 |
| pLR24 | Yep_URA17_pGAL1-3Xflag-M60TASYBRE-tCYC1 |
| pLR25 | Yep_Leu2d_pGAL1-3Xflag-M60TASYBRE-tCYC1 |
| pLR33 | Yep_pTDH3-M60TASYBACA-tCYC1 |
| pLR34 | Yep_pTDH3-M50TTASYBRE-tCYC1 |
| pLR35 | Yep_pTDH3-KOZ-M50TTASYBRE-tCYC1 |
| pLR36 | Yep_pTDH3--M60TASYBRE-MBP-tADH1<br>pGAL1-T5aH-tCYC1 |
| pLR37 | Yep_pTDH3-CAS9-tADH1 |
| pLR40 | Yep_pGAL1-M40TASYBRE-tCYC1 |
| pLR41 | Yep_pGAL1-M44TASYBRE-tCYC1 |
| pLR42 | Yep_pGAL1-M48TASYBRE-tCYC1 |
| pLR43 | Yep_pGAL1-M52TASYBRE-tCYC1 |
| pLR44 | Yep_ pGAL1-M54TASYBRE-tCYC1 |
| pLR45 | Yep_ pGAL1-M56TASYBRE-tCYC1 |

| | |
|---|---|
| pLR46 | Yep_ pGAL1-M58TASYBRE-tCYC1 |
| pLR47 | Yep_ pGAL1-M62TASYBRE-tCYC1 |
| pLR48 | Yep_ pGAL1-M64TASYBRE-tCYC1 |
| pLR49 | Yep_ pGAL1-M66TASYBRE-tCYC1 |
| pLR50 | Yep_ pGAL1-M68TASYBRE-tCYC1 |
| pLR51 | Yep_ pGAL1-M70TASYBRE-tCYC1 |

Supplementary Table 2: **List of primers used in the study**

| **Name** | **Sequence** |
|---|---|
| M13 F Backbone Ups of Promoter | GTAAAACGACGGCCAGT |
| M13 R backbone down of Term | CAGGAAACAGCTATGAC |
| TASY1 | GGTAGAGACATGTTGGCCCA |
| TASY2 | AGCGGAGTATTCTGGTTCGA |
| TASY3 | TCCGTCATTGCCTTGTCTGT |
| TASY4 | ACCGATACCCAAATGTTCAATATTGT |
| beginning of TASY with 3Xflag F | TGACAAATCCTCTTCCACTGGT |
| 213 BP TASY R | ATCACCCAAGGCGTTGAACA |
| TASY5 | TCTGCTGGTCAAACTCACGT |
| Backbone R after Term | CAAGCGCGCAATTAACCCTC |
| (backbone1)_forward | TACAGACACTGGTCTGAACGTGGTATTGGTTGGGG |

| | |
|---|---|
| (backbone1)_reverse | GCGGCCAGCAAAACTAAGTTTCTTAGTATGATCCAATATCAAAGGAAATG |
| (URA17)_forward | TGGATCATACTAAGAAACTTAGTTTTGCTGGCCGCATCTTCTCAAATATGC |
| (ura17)_reverse | AGACGAAACTATATACGCAGGAAACGAAGATAAATCATGTCGAAA |
| (bakbone2)_forward | CGACATGATTTATCTTCGTTTCCTGCGTATATAGTTTCGTCTACCCTATGAACA |
| (Backbone2)_reverse | ATACCACGTTCAGACCAGTGTCTGTAGACGTAATCCAAAGCACC |
| (PArt1TASY)_forward | AAGCTCCCTCGTGCGCTCTCCTGTTCCGACC |
| (PArt1TASY)_reverse | AGAAATATCTTGACCGCAGTTTATATATATTTCAAGGATATACCATTCTAA TGT CTG |
| (part2TASY)_forward | CCTTGAAATATATATAAACTGCGGTCAAGATATTTCTTGAATCAGGCGC |
| (part2TASY)_reverse | GAACAGGAGAGCGCACGAGGGAGCTTCCAGGGGG |
| (Backbone1)_forward | GTTTTAGAGCTAGAAATAGCAAGTTAAAATAAGGC |
| (Backbone1)_reverse | GACATAACTAATTACATGACTCGAGAAGAGATCACACCTTCCT |
| (Backbone2)_forward | AGGTGTGATCTCTTCTCGAGTCATGTAATTAGTTATGTCACGCTTACGTTCACGCCC |
| (Backbone2)_(CAN1Y)_reverse | AGCCTTATTTTAACTTGCTATTTCTAGCTCTAAAACTCCTCCATAGAGAACGTATCAAAGTCCCATTCGCCACCCGAAGG |
| Gal1p F 3/4 | GGTAATTAATCAGCGAAGCGATG |
| Gal1p F 1/2 | CCCACAAACCTTCAAATCAACG |
| 3xflag3 | GGTTTGAATGACGTTATGATGCC |

| | |
|---|---|
| 3Xflag4 | CAAGGCGTTGAACATATCCTTG |
| DOWNTASY and UPCYC1 R | TCGGTTAGAGCGGATTCAAACT |
| Backbone after CYC1T R | CGC AAC GCA ATT AAT GTG AGT |
| (ORICYC1)_(KOZAK-SEQ)_reverse | CCGCAGACATTTTTTTGTTTGTTTATGTGTGTTTATTCGAAAC |
| GAL1 1/2 F | AGTAACCTGGCCCCACAAAC |
| Reverse3Xflag | ACCCAATTTAAGGCTTGTGGGA |
| (ORICYC1)_forward | GGTATATCCTTGAAATATATATAGGAGAAAATACCGCATCAGGAAATTG |
| BACKBONEtruncatedleu2d)_reverse | CCTGATGCGGTATTTTCTCCTATATATATTTCAAGGATATACCATTCTAATGTCTGCCCCTAAGAAGATCGTCG |
| CYC1T F Beginning | ATCCGCTCTAACCGAAAAGGAAGG |
| (vectorpart1)_reverse CASURA17 | GATTTTGTTGATGCCATTATAGTTTTTTCTCCTTGACGTTAAAGTATAGAGG |
| (vectorpart1)_forward CAS URA17 | GGTGTCTATTTTCTCTTCCATAAAAAAAGCCTGACTCCACTTCCCGC |
| vectorpart2_forward CASURA17 | TTAATTTGCGGCCAAGCTTGATATCGAATTCCTGCAGCCCG |
| vectorpart2_reverse CASURA17 | TGGAGTCAGGCTTTTTTTATGGAAGAGAAAATAGACACCAAAGTAGCC |
| term backbone kozak reverse | GAAAATCCTTGCTTAATCATCACCGAAACGCGCG |
| term backbone kozak forward | CCAATTCAAGTTTGAATCCGCTCTAACCGAAAAGGAAGG |

| | |
|---|---|
| TASYN50 kozak reverse | CCTTTTCGGTTAGAGCGGATTCAAACTTGAATTGGATCAATGTAGAC |
| TASYN50 kozak forward | CTGGTCCTGTCGTAATGTCCTCTTCCACTGGTACCTC |
| promoter backbone kozak reverse | GGAAGAGGACATTACGACAGGACCAGGACCACCACTTCCTCTCATAGACATTTTTTTGTTTGTTTATGTGTGTTTATTCGAAAC |
| promoter backbone kozak forward | CGTTTCGGTGATGATTAAGCAAGGATTTTCTTAACTTCTTCGGC |
| term backbone reverse | GAAAATCCTTGCTTAATCATCACCGAAACGCGCG |
| term backbone forward | CCAATTCAAGTTTGAATCCGCTCTAACCGAAAAGGAAGG |
| TASYN50 reverse | CCTTTTCGGTTAGAGCGGATTCAAACTTGAATTGGATCAATGTAGAC |
| TASYN50 forward | CTGGTCCTGTCGTAATGTCCTCTTCCACTGGTACCTC |
| promoter backbone reverse | GGAAGAGGACATTACGACAGGACCAGGACCACCACTTCCTCTCATTTTGTTTGTTTATGTGTGTTTATTCGAAAC |
| promoter backbone forward | CGTTTCGGTGATGATTAAGCAAGGATTTTCTTAACTTCTTCGGC |
| Beginning- TASY MBP primer F | TCCAGTTCGAGTTTATCATTATCAATACTGCCA |
| Beginning-TASY MBP primer R | GGTTTCAGAAACGACCTTAGAGGTACCAGTGGAAGAGG |
| End- TASY MBP primer F | GGTACCTCTAAGGTCGTTTCTGAAACCTCCTCTACCATCGTCG |
| End- TASY MBP primer R | GGCTTCTAATCCGTACTGGAGTTAGCATATCTACAATTGGGTGAAATGGGG |
| CRISPR Backbone 1 forward | GTTTTAGAGCTAGAAATAGCAAGTTAAAAT |

| | |
|---|---|
| CRISPR Backbone 1 reverse | TGGAGAATATACTAAGGGTACTGTTGACATTGCGAAGAGCGACAAAGATT |
| CRISPR Backbone 2 forward | CCGATAACAAAATCTTTGTCGCTCTTCGCAATGTCAACAGTACCCTTAGTATATTCTCCA |
| CRISPR Backbone 2 reverse | AAAGTCCCATTCGCCACCCGAA |
| (upstream)_reverse | CTAGCTCTAAAACTCTCCCGGGGGCGAGTCG |
| (upstream)_forward | CGTTTACAATTTCCTGATGCGGTATTTTCTCCTTACGC |
| F-up1622b GAL1 | AACATTTAAGTCACAAGGAGGAATATCAGTT |
| R-up1622b | GCAGTATTGATAATGATAAACTCGAACTGAACTACTTTTCTTAAACTGTCAACAGCCA |
| F-dn1622b | TCACCCAATTGTAGATATGCTAACTCCGTAGATACTCGTCTTACGAAATTGGATATAGTT |
| R-dn1622b | ACTTTGGAAAAGAAGGTACGGACTACT |
| F-up1014a GAL1 | TATTGACCAGTAGTCATATTACTGGCATATTATC |
| R-up1014a | GCAGTATTGATAATGATAAACTCGAACTGAGGATATTAATTTTAGGGTCTCTTGATGCAC |
| F-dn1014a | ACCCAATTGTAGATATGCTAACTCCCACTGTTTTCATCTAGACGTGGGAC |
| R-dn1014a | GCAATACCAGAGATGACTGGCC |
| F-TDH3-TASY-MBP-ADH1 | CAGTTCGAGTTTATCATTATCAATACTGCC |
| R-TDH3-TASY-MBP-ADH1 | GGAGTTAGCATATCTACAATTGGGTGAA |
| ADH1 fwd | CCCAAGACCATAAGCGAATTTCTTATG |
| ADH1 rev | GGAATTGTGAGCGGATAACGGAGTTAGCATATCTAC |

| | |
|---|---|
| 1622bUP 8/10 F | TGCGAGCAGACTTTGTCCAT |
| 1622bDOWN 2/10 R | TCGTTCTACAAGTCCAGCCA |
| 1014a UP 8/10 F | AGGATTTCTATGTTCTCGAGGAGA |
| 1014a DOWN 2/10 R | TCCGCCCTTTGCATCTATAAA |
| 1014 (Scaffold)_forward | TTATGTGCGTATTGCTTTCAGTTTTAGAGCTAGAAATAGCAAGT TAAAATAAGGC |
| Scaffold_reverse | GCACCGACTCGGTGCCAC |
| Scaffold forward | GTTTTAGAGCTAGAAATAGCAAGTTAAAATAAGGC |
| (1622b)_(Scaffold)_forward | GTCACGTTCCTGAGGTTACTGTTTTAGAGCTAGAAATAGCAAGT TAAAATAAGGC |

Supplementary Table 3: **DNA sequences**

| **Gene** | **Sequence** | **Reference** |
|---|---|---|
| *E. coli MBP* (Yeast codon-optimised) | TCTAAGATTGAAGAAGGTAAGTTGGTTATCTGGAT TAACGGTGACAAGGGTTACAACGGTTTGGCTGAAG TTGGTAAGAAATTTGAAAAAGATACCGGTATCAAG GTCACTGTTGAACACCCAGACAAGTTGGAAGAAA AGTTTCCACAAGTTGCTGCCACTGGTGATGGTCCA GACATTATCTTCTGGGCTCATGACAGATTCGGTGG TTACGCCCAATCCGGTTTGTTAGCCGAGATCACCC CAGATAAGGCTTTTCAAGATAAGTTGTATCCATTC ACTTGGGATGCCGTCAGATACAACGGTAAGTTAAT CGCCTACCCAATTGCTGTTGAAGCTTTGTCTTTGAT CTACAATAAGGACTTGTTACCTAACCCACCAAAGA | (Sun, Tropea and Waugh, 2011) |

| | | |
|---|---|---|
| | CCTGGGAAGAAATCCCAGCTTTAGATAAGGAGTTA<br>AAAGCTAAGGGTAAGTCCGCTTTGATGTTTAACTT<br>GCAAGAACCATACTTCACTTGGCCATTGATCGCTG<br>CTGATGGTGGTTACGCTTTTAAGTATGAAAACGGT<br>AAATACGACATTAAGGATGTCGGTGTCGACAATGC<br>TGGTGCTAAGGCCGGTTTAACTTTCTTAGTCGATTT<br>GATTAAGAATAAACATATGAATGCTGACACTGATT<br>ACTCTATTGCTGAAGCTGCTTTCAACAAGGGTGAA<br>ACCGCTATGACTATTAACGGTCCATGGGCCTGGTC<br>TAACATTGATACCTCTAAAGTCAACTACGGTGTCA<br>CCGTCTTGCCAACTTTTAAGGGTCAACCATCTAAG<br>CCATTCGTCGGTGTCTTGTCTGCCGGTATTAACGCT<br>GCCTCTCCAAATAAGGAATTGGCCAAGGAATTCTT<br>AGAAAACTACTTGTTAACCGATGAAGGTTTAGAGG<br>CCGTTAACAAGGATAAGCCATTAGGTGCTGTTGCT<br>TTGAAGTCTTACGAAGAAGAGTTGGCTAAGGATCC<br>AAGAATTGCTGCTACTATGGAAAACGCTCAAAAG<br>GGTGAAATTATGCCAAACATCCCACAAATGTCTGC<br>TTTCTGGTACGCTGTTCGTACCGCCGTCATTAATGC<br>CGCTTCTGGTCGTCAAACTGTTGATGAAGCCTTGA<br>AGGACGCTCAAACCAGAATTACTAAG | |
| Mutant variant of *Aequorea victoria GFP* | AGTAAAGGAGAAGAACTTTTCACTGGAGTTGTCCC<br>AATTCTTGTTGAATTAGATGGTGATGTTAATGGGC<br>ACAAATTTTCTGTCAGTGGAGAGGGTGAAGGTGAT<br>GCAACATACGGAAAACTTACCCTTAAATTTATTTG | (Houser *et al.*, 2012) |

| | |
|---|---|
| | CACTACTGGAAAACTACCTGTTCCATGGCCAACAC<br>TTGTCACTACTCTCACTTATGGTGTTCAATGCTTTT<br>CAAGATACCCAGATCAcATGAAACaGCATGACTTTT<br>TCAAGAGTGCCATGCCCGAAGGTTATGTACAGGAA<br>AGAACTATATTTTTCAAAGATGACGGGAACTACA<br>AGACACGTGCTGAAGTCAAGTTTGAAGGTGAT<br>ACCCTTGTTAATAGAATCGAGTTAAAAGGTAT<br>TGATTTTAAAGAAGATGGAAACATTCTTGGAC<br>ACAAATTGGAATACAACTATAACTCACACAAT<br>GTATACATCATGGCAGACAAACAAAAGAATG<br>GAATCAAAGcTAACTTCAAAATTAGACACAAC<br>ATTGAAGATGGAAGCGTTCAACTAGCAGACCA<br>TTATCAACAAAATACTCCAATTGGCGATGGCC<br>CTGTCCTTTTACCAGACAACCATTACCTGTCCA<br>CACAATCTGCCCTTTCGAAAGATCCCAACGAA<br>AAGAGAGACCACATGGTCCTTCTTGAGTTTGT<br>AACAGCTGCTGGGATTACACATGGCATGGATG<br>AACTATACAAA |
| Yeast codon-optimised *TASY* | ATGTCCTCTTCCACTGGTACCTCTAAGGTCGTTTCT<br>GAAACCTCCTCTACCATCGTCGATGACATTCCAAG<br>ATTGTCTGCTAATTACCACGGTGACTTGTGGCATC<br>ATAACGTCATTCAAACCTTGGAAACTCCATTTAGA<br>GAATCTTCTACTTATCAAGAGAGAGCTGATGAATT<br>GGTTGTCAAGATCAAGGATATGTTCAACGCCTTGG |

| | |
|---|---|
| | GTGATGGTGATATCTCTCCATCTGCTTATGATACTG<br>CCTGGGTCGCTAGATTGGCTACCATCTCTTCCGAC<br>GGTTCCGAAAAGCCAAGATTCCCACAAGCCTTAAA<br>TTGGGTTTTTAACAACCAATTGCAAGACGGTTCTT<br>GGGGTATTGAATCTCATTTCTCTTTGTGTGATAGAT<br>TGTTGAACACCACTAACTCCGTCATTGCCTTGTCTG<br>TTTGGAAGACTGGTCACTCTCAAGTTCAACAAGGT<br>GCCGAATTCATTGCCGAAAACTTGAGATTATTGAA<br>CGAAGAAGATGAATTGTCTCCAGACTTCCAAATCA<br>TTTTTCCAGCTTTGTTGCAAAAGGCCAAGGCCTTA<br>GGTATCAACTTGCCATACGACTTGCCATTCATCAA<br>GTACTTGTCTACTACCAGAGAAGCTAGATTGACTG<br>ACGTCTCCGCTGCTGCTGACAACATTCCAGCCAAC<br>ATGTTGAATGCCTTGGAAGGTTTAGAAGAAGTCAT<br>TGATTGGAACAAGATTATGAGATTCCAATCTAAAG<br>ACGGTTCTTTTTTGTCTTCCCCTGCTTCTACTGCTTG<br>TGTCTTGATGAACACCGGTGATGAGAAGTGTTTCA<br>CTTTCTTGAATAACTTGTTGGATAAATTCGGTGGTT<br>GTGTTCCATGTATGTATTCCATTGATTTATTGGAAA<br>GATTGTCTTTAGTTGACAATATTGAACATTTGGGT<br>ATCGGTAGACACTTCAAGCAAGAAATTAAGGGTG<br>CTTTGGATTACGTCTACAGACACTGGTCTGAACGT<br>GGTATTGGTTGGGGTAGAGATTCTTTGGTTCCAGA<br>TTTAAACACTACTGCCTTGGGTTTGCGTACCTTGAG<br>AATGCACGGTTACAACGTTTCTTCCGACGTTTTGA |

| | |
|---|---|
| | ACAACTTCAAGGATGAAAACGGTAGATTCTTTTCC<br>TCTGCTGGTCAAACTCACGTCGAATTAAGATCCGT<br>TGTCAACTTGTTCAGAGCTTCTGATTTGGCCTTCCC<br>AGACGAAAGAGCTATGGATGATGCTAGAAAATTC<br>GCTGAACCATATTTGAGAGAGGCCTTGGCCACCAA<br>GATTTCTACCAACACTAAGTTGTTCAAGGAGATTG<br>AATACGTTGTCGAATACCCATGGCACATGTCCATC<br>CCAAGATTGGAAGCTAGATCCTATATTGACTCCTA<br>CGACGATAACTACGTTTGGCAAAGAAAAACCTTGT<br>ACCGTATGCCATCCTTGTCTAACTCCAAGTGTTTGG<br>AGTTAGCTAAATTAGACTTCAATATCGTCCAATCC<br>TTACATCAAGAAGAATTGAAATTGTTGACCAGATG<br>GTGGAAGGAATCTGGTATGGCTGATATCAACTTCA<br>CCAGACACAGAGTCGCCGAAGTTTACTTCTCCTCT<br>GCTACTTTCGAACCAGAATACTCCGCTACCAGAAT<br>TGCTTTCACTAAGATCGGTTGTTTACAAGTTTTATT<br>CGATGATATGGCTGACATTTTCGCTACTTTGGATG<br>AATTGAAGTCTTTCACTGAAGGTGTTAAGAGATGG<br>GATACTTCTTTGTTGCATGAAATCCCAGAATGTAT<br>GCAAACCTGTTTTAAGGTTTGGTTTAAGTTGATGG<br>AGGAAGTCAACAACGACGTTGTTAAGGTTCAAGGT<br>AGAGACATGTTGGCCCACATCAGAAAGCCATGGG<br>AATTGTACTTCAACTGTTACGTCCAAGAAAGAGAA<br>TGGTTGGAAGCTGGTTATATCCCTACCTTCGAAGA<br>ATACTTGAAGACTTACGCTATCTCTGTCGGTTTAG |

| | |
|---|---|
| | GTCCTTGTACTTTGCAACCAATTTTGTTGATGGGTG<br>AATTAGTTAAGGACGATGTTGTTGAAAAAGTTCAC<br>TACCCATCCAACATGTTCGAATTAGTTTCCTTGTCC<br>TGGAGATTAACCAATGACACCAAGACTTACCAAGC<br>CGAAAAGGCTAGAGGTCAACAAGCTTCTGGTATCG<br>CTTGTTACATGAAGGATAATCCAGGTGCCACTGAA<br>GAAGATGCTATTAAGCACATTTGTAGAGTTGTTGA<br>CAGAGCTTTGAAGGAAGCCTCTTTCGAATACTTTA<br>AGCCATCCAACGATATCCCAATGGGTTGCAAGTCT<br>TTCATCTTCAACTTGAGATTGTGTGTCCAAATCTTC<br>TACAAGTTTATCGACGGTTACGGTATTGCTAACGA<br>AGAAATCAAGGATTACATTAGAAAAGTCTACATTG<br>ATCCAATTCAAGTTTGA |

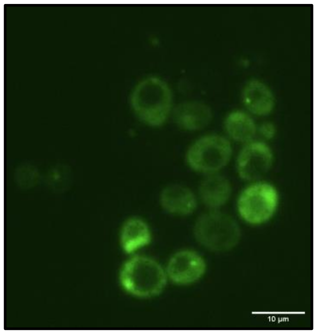

Supplementary Figure 1: **GFP tag attached to *TASY* show spotted subcellular localization consistent with poor TASY solubility as imaged by confocal fluorescence microscopy.**

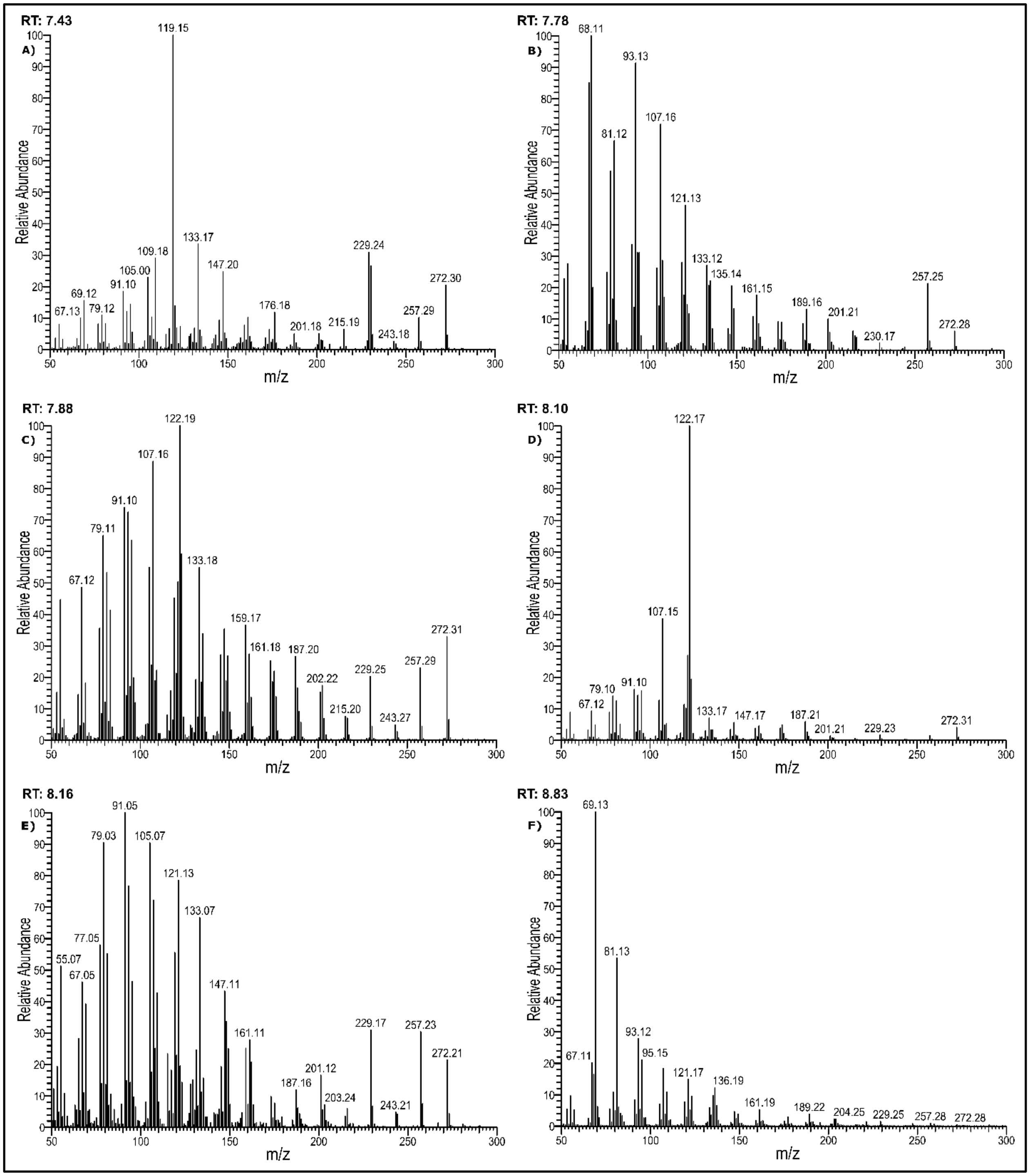

Supplementary Figure 2: **Mass spectra for compounds produced by LRS5**. A) Verticillene; B) Diterpene 1; C) Iso-taxadiene; D) Taxadiene; E) Diterpene 2; F) GGOH

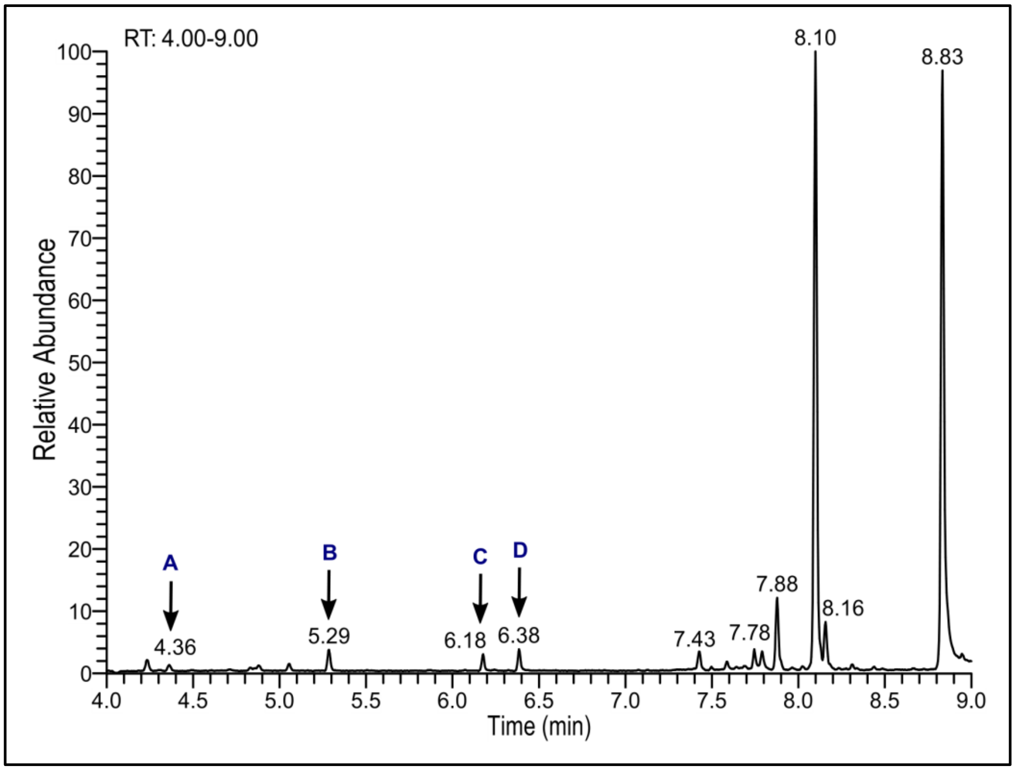

Supplementary Figure 3: **LRS5 gas chromatogram showing additional potential terpenoids.** Compounds produced during LRS5 shake flask cultivation at 30 °C. The mass spectra of unknow potential terpenoids A, B, C and D are shown in the Supplementary Figure 4.

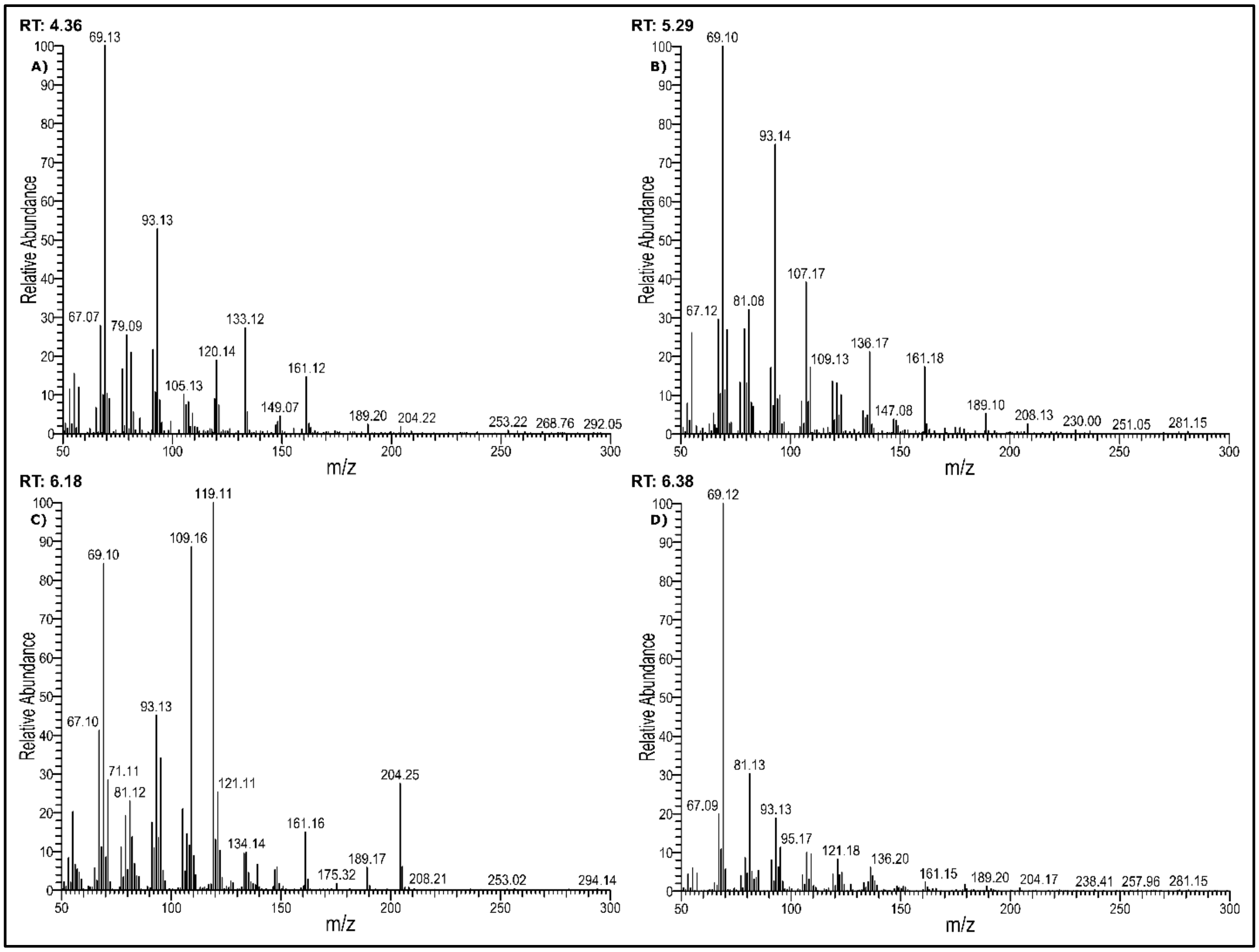

Supplementary Figure 4: **Mass spectra of the additional potential terpenoids (A-D) produced by LRS5 and shown in the chromatogram in Supplementary Figure 3.**
